# Spatial factors predict variation in reports of human-wildlife interactions but not public attitudes towards a widespread urban carnivore, the red fox

**DOI:** 10.1101/2025.07.23.666373

**Authors:** Kristy A. Adaway, Charlotte R. Hopkins, Carl D. Soulsbury, F. Blake Morton

**Author notes:** Corresponding author: Kristy Adaway, Department of Psychology, University of Hull, HU6 7RX, United Kingdom.

## Abstract

Despite growing recognition that spatial factors such as urbanisation and geographic region shape human-wildlife interactions, few studies have examined – on a large geographic scale – how spatial factors reflect the subjective human dimensions to those interactions, including public attitudes and self-reported encounters with wildlife. Understanding how spatial factors shape these dimensions may have implications for urban rewilding because they can reflect local tolerance of animals and may potentially predict human-wildlife interactions. We examined how urbanisation type and geographic region are related to people’s attitudes, reported encounters (public feeding and bin-raiding), and reported use of control measures towards the world’s most urbanised terrestrial carnivore, the red fox, *Vulpes vulpes*. Online survey data were obtained from 1,275 participants in the United Kingdom. Reports of fox bin-raiding were associated with increased reports of wildlife feeding, more negative attitudes towards foxes, and greater reported use of professional and do-it-yourself control measures. However, the role of spatial factors varied among these relationships. Specifically, urbanisation and geographic region significantly predicted reports of fox bin-raiding, with urban residents – particularly from London – more likely to report such behaviour. Urbanisation predicted reports of wildlife feeding and use of professional pest control of foxes within their area. Geographic region predicted reports of do-it-yourself control measures. Neither urbanisation nor geographic region significantly predicted public attitudes towards foxes. Together, these findings highlight the complex interactions between spatial context and the subjective human dimensions of wildlife interactions, underscoring the importance of nuanced, context-specific strategies to support human-wildlife coexistence and urban rewilding initiatives.

**Highlights:**

- Urban rewilding success depends on public tolerance of local wildlife.
- We studied spatial factors and subjective human dimensions of fox interactions.
- Reports of bin-raiding, wildlife feeding, fox-related attitudes, and pest control covaried.
- The role of urbanisation and geography varied in predicting these reports.
- Such spatial complexity should be considered in designing urban rewilding strategies.

## Introduction

Urbanisation is currently one of the fastest forms of human-driven landscape change on the planet (Li et al., 2022; Simkin et al., 2022). As the world’s global human population continues to rise, 68% of the global population are predicted to live in urban areas by 2050 (Li et al., 2022; Simkin et al., 2022; United Nations, 2019). Urban environments create challenging environments for wildlife to inhabit, including high human population density, road density, the loss and fragmentation of natural habitat, pollution, and invasive species (McKinney, 2002, 2006; Ramadhan, 2024). Such ecological impacts from urbanisation highlight the urgent need to understand the changing relationship between humans and the natural world (Cox, Hudson, et al., 2017; Gaston & Soga, 2020; Soga & Gaston, 2016).

Species richness of wildlife in cities varies across urban-rural gradients, but the most heavily urbanised areas often show a lower number of species (Aronson et al., 2014; McKinney, 2002, 2006; Ramadhan, 2024; Sidemo-Holm et al., 2022; Sol et al., 2014). Thus, urbanisation typically reduces access and diversity in opportunities for people to physically, cognitively, and emotionally “connect” with nature (Cox & Gaston, 2018; McEwan et al., 2019; Richardson et al., 2022; Soga & Gaston, 2016). Indeed, research shows a global decline in nature connectedness (Cazalis et al., 2023; Soga & Gaston, 2016), and urban residents often experience lower levels of nature connectedness compared to rural residents (Barragan-Jason et al., 2025; Kageyama et al., 2024; Richardson et al., 2022). Declining human-nature connectedness can lead to poorer mental and physical health, such as depression and anxiety (Barrable & Booth, 2022; Cox, Shanahan, et al., 2017; Martyn & Brymer, 2016; Sibthorpe & Brymer, 2020; White et al., 2021). It can also, at least in some instances, reduce proenvironmental attitudes and behaviour (Barrable & Booth, 2022; Lengieza et al., 2023; Otto & Pensini, 2017; Straka et al., 2025; Whitburn et al., 2020), shifting people’s priorities away from engaging with nature, including support for conservation initiatives (Barrows et al., 2022; Richardson et al., 2020). Thus, to help foster deeper connections to nature and support for biological conservation efforts, it is crucial that researchers understand how to improve people’s experiences within nature, especially in cities (Cox & Gaston, 2018; Richardson et al., 2022; Soga & Gaston, 2016).

“Rewilding”, the process of restoring key natural processes and missing components of the food web to create self-sustaining ecosystems (Carver et al., 2021), has been suggested as a way to increase people’s access to nature (Lengieza et al., 2025). More specifically, “urban rewilding”, defined broadly as low management initiatives that seek to improve biodiversity in urban environments (Pettorelli et al., 2022), has been proposed as a way to improve access to nature within cities (Finnerty et al., 2025; Pettorelli et al., 2022; Webb & Moxon, 2023). Urban rewilding initiatives can include the installation of green roofs to buildings or the establishment of wildflower meadows to provide valuable habitat for urban wildlife (Griffiths-Lee et al., 2022; Marshall et al., 2023; Wooster et al., 2022). Similarly, adding wildlife-friendly features in gardens, such as ponds and native vegetation, can make space for nature by offering resources and refuge to various species (Hill et al., 2017; Threlfall et al., 2017). Using these strategies to encourage more wildlife to urban areas and gardens can improve nature connectedness and well-being by providing residents with greater opportunities to experience positive interactions with wildlife. For instance, bird watching in residents’ own back gardens for a short period of time (e.g., 30 minutes) can foster deeper connections with nature, promoting better health and well-being in those individuals (White et al., 2023). Given the benefits of urban rewilding, these initiatives are now being actively promoted and implemented in many cities across the world, including major cities such as London, New York, Singapore, and Sydney (Arup & C40 Cities, 2022; Finnerty et al., 2025). However, while these initiatives aim to foster more positive relationships between people and nature, it remains critical to monitor human-wildlife interactions to guide effective management of them.

Public reports of human-wildlife interactions, collected through anecdotes, interviews, or self-reported questionnaires, are often the primary source of information on human-wildlife interactions (Moesch, Jeschke, et al., 2024; Morton & Soulsbury, 2025). Such reports, however, are subjective and depend on public perceptions, including what people think they experience (e.g., types and frequencies of interactions), and in terms of public attitudes, including how people evaluate those perceived interactions (e.g., positive versus negative emotions) (Morton & Soulsbury, 2025; Nyhus, 2016). Public perceptions and attitudes likely play an important role in shaping the success of conservation initiatives, including urban rewilding, such as in terms of encouraging behavioural changes within communities (Bruskotter & Fulton, 2012; Bruskotter & Wilson, 2014; Conejero et al., 2019; K.J. Hobson et al., 2024; Naughton-Treves et al., 2003; Puri et al., 2024). Indeed, socio-psychological research shows that people are unlikely to change their behaviour unless it aligns specifically with what they already know, value, or believe (Toomey, 2023). Thus, in terms of urban rewilding, for instance, encouraging people to adopt pro-environmental behaviours, such as planting natural food for wildlife within gardens, is likely to be more effective when campaigns are informed by residents’ pre-existing perceptions and attitudes about those animal species.

Despite growing recognition that spatial factors such as urbanisation and geographic region are associated with differences in human-wildlife interactions, few studies have examined – on a large geographic scale – how spatial factors are related to the subjective human dimensions to these interactions, including self-reported encounters, attitudes, and behaviour towards wildlife within the same study system (Baker et al., 2020; Bruskotter et al., 2015; Bruskotter & Wilson, 2014; Dunn et al., 2018; Keziah J Hobson et al., 2024; Kansky et al., 2014; Puri et al., 2024). Research has shown, for example, that attitudes related to tolerance of wildlife varies across urban-rural gradients (Balčiauskas & Kazlauskas, 2014; Bjerke et al., 1998; Kansky et al., 2014; Kimmig et al., 2020; Ostermann-Miyashita et al., 2023; Tan et al., 2020), with some urban residents reporting greater acceptance of wildlife due to more frequent exposure (Moesch, Straka, et al., 2024). However, such patterns are not universal and may depend on other factors, such as nuisance animal behaviours (e.g., bin-raiding) and other perceived or actual risks (e.g., disease and physical attacks) (Kansky et al., 2014; König et al., 2020; Tan et al., 2020). Thus, understanding spatial variation in human dimensions is important to ensure that conservation, including urban rewilding, is locally-tailored to help monitor human-wildlife interactions across different areas (Soulsbury & White, 2015).

We examined how urbanisation type (urban, suburban, or rural) and geographic region are related to people’s reports of bin-raiding, food provisioning habits, attitudes, and reported use of control measures towards the world’s most urbanised terrestrial carnivore, the red fox, *Vulpes vulpes*. Red foxes (hereafter foxes) are an ideal case study because they are the most globally widespread terrestrial urban carnivore on the planet (Bijl & Csányi, 2022; Castañeda et al., 2022; Soe et al., 2017; Soulsbury et al., 2010) and have a long and complex history of living within urban areas across many countries (Baker et al., 2006; Basak et al., 2022; Bridge & Harris, 2020; Contesse et al., 2004; Moesch, Jeschke, et al., 2024). Although human-fox interactions are frequent and varied, the current study specifically chose to investigate spatial variation in food provisioning by residents and bin-raiding by foxes because they are two of the most commonly-reported interactions with foxes within neighbourhoods (Baker et al., 2020; Brand & Baldwin, 2020; Harris, 1981; Scott et al., 2014). Because foxes are often more active within and around urban areas compared to the countryside (Scott et al., 2014; Morton et al., 2023), urban residents are potentially more likely to feed them (Baker et al., 2004), which may also lead to a higher prevalence of reported conflict with these animals, including bin-raiding to exploit human food (Brand & Baldwin, 2020).

Reports of human-fox conflicts, including bin-raiding, have historically reduced public tolerance of foxes, leading some people to advocate for greater use of pest control to manage their populations (Baker et al., 2020; Brand & Baldwin, 2020; Kimmig et al., 2020). Currently, measures of wildlife control are commonly employed in many urban areas for species labelled as “pests”, including foxes (Baker et al., 2020; McCleery et al., 2012; McPherson et al., 2021; Sato, 2017; Schell et al., 2021; Vantassel & Groepper, 2015). While some wildlife control measures may be necessary when public health or safety are put at risk (Baker et al., 2020; Puri et al., 2024), in most other cases, they can be avoided through simple preventative measures, such as building fences and other barriers to deter animals from entering an area (The_Humane_Society_of_the_United_States, 2023).

The current study hypothesised that individuals who feed wildlife would be more likely to report seeing foxes raid bins within their neighbourhood. We also hypothesised that people who reported fox bin-raiding behaviour – regardless of whether or not they fed wildlife – would also report more negative attitudes and greater use of fox control measures. Finally, to understand how these patterns varied across different spatial contexts, we assessed whether urbanisation type and geographic region explained differences in people’s reports of wildlife feeding, fox bin-raiding, public attitudes towards foxes, and the use of fox control measures.

## Methods and materials

### Ethical statement

The study was approved by the Faculty of Health Sciences Ethics Committee of the University of Hull (FHS 22-23.37) and followed the research guidelines of the British Psychological Society.

### Public attitudes towards foxes

A nation-wide online survey assessing public attitudes about foxes was conducted in Great Britain between March and April 2023. Participants were recruited using a commercial survey company (Survation) to ensure a representative sample using the company’s existing participant pool. The survey was originally conducted as part of a study that investigated whether communicating information about the psychology of foxes influenced public attitudes towards the species, and whether this was conditional on how that information was disseminated (i.e., press release or a video) (Morton et al., 2024). Participants’ attitudes towards foxes were not significantly influenced by the information or format given to them (Morton et al., 2024), and so we used these attitude scores in the current study. A total of 1,373 people participated in the study, with an average age of 53 ± 18.2 years (range: 18-93 years) and 689 and 684 people identifying as male or female, respectively. Online surveys allow rapid data collection and access to large, geographically dispersed samples, but they also have potential limitations, such as self-selection bias. Nevertheless, as discussed earlier, subjective reports of human-wildlife interactions, including data collected through surveys, are often the primary source of information on human-wildlife interactions and, hence, play an important role in understanding the subjective human dimensions to human-wildlife interactions.

Full details of the survey can be found in Morton et al. (2024). We used data from the final section of that survey, where participants were asked to what extent they agree with 24 statements made about foxes on a 7-point Likert scale (e.g., “I enjoy seeing foxes”). These responses were then used to measure their attitudes towards foxes. Participants’ responses were analysed using a Principal Component Analysis (PCA), which suggested that the data were comprised of a single component characterised by people’s attitudes towards the species. Thus, an overall attitude score was determined for each participant by calculating the average score for questions describing positive attitudes, and the average score for questions that described negative attitudes, and then subtracting the negative from the positive. Participants’ scores ranged from −6 (very negative) to 6 (very positive) and were classified as being relatively more positive than negative in terms of their overall attitude towards foxes if their score was greater than 0.

### Urbanisation type, geographic region, and subjective reports of human-fox interactions

Data on participants’ geographic region, level of urbanisation, and subjective reports of human-fox interactions from Morton et al. (2024) were used. We used the first half of participants’ postcode to categorise participants into 11 regions, including East Midlands, East of England, London, North East, North West, Scotland, South East, South West, Wales, West Midlands, and Yorkshire and the Humber. Urbanisation type was based on three factors, including urban (i.e., a town or city, lots of people live there, and there are lots of different kinds of buildings close together), suburban (i.e., mostly houses, located just outside a city or town), or rural (i.e., mostly countryside with few houses or buildings). Since there is no single best method to define urbanisation types, we operationalised our definitions for urban, suburban, rural classifications following Morton and Soulsbury (2025), where we asked participants to evaluate their physical surroundings based on broad characteristics, ranging from mostly countryside (rural) to having lots of people/houses close by (urban). In the UK, towns and cities have official designated boundaries on maps and roads, and common terminology distinguishes urban, suburban, and rural areas using the same or similar descriptions used in the current study. While we would not expect perfect agreement between self-reported and satellite-derived measures of people’s urban-rural environment, studies do show that people’s subjective evaluations of urban, suburban, and rural classifications are broadly reflective of more objective geospatial measures (Gianotti et al., 2016).

To assess reports of human-fox interactions among participants, two key variables were derived. First, participants were asked whether they had ever seen a fox eating rubbish from their outdoor bin, or from a public bin owned by their local council. Responses to these two questions were combined into a single binary measure of prior experience with local fox bin-raiding behaviour, classified as yes (1) if they had personally seen a fox raiding either type of bin, or no (0) if they had not seen a fox raiding a bin. Second, participants were asked whether they ever leave food outside for foxes, or for other wildlife (e.g., birds or hedgehogs). Given the relatively small number of participants who fed only foxes (N = 8 people) (Morton et al., 2024), participants’ food provisioning activities were based on three factors, including people who fed all wildlife (foxes and other species), people who only fed other wildlife (not foxes), and people who did not feed any wildlife. To further understand the potential impact of observing fox bin-raiding behaviour on people’s tolerance (in addition to evaluating their attitudes), participants were also asked whether they had ever contacted a pest control service to resolve an issue they had with foxes (hereafter “professional services”), and whether they had ever tried to harm or kill a fox themselves (hereafter “do-it-yourself” or DIY measures), at their place of residence. We analysed professional services and DIY measures separately, rather than as a combined ‘pest control’ score, because they potentially reflect different underlying factors. For example, people’s willingness to use professional services may be influenced by socioeconomic status or access to those services, whereas people’s willingness to engage in DIY could reflect differences in aversion to risk or disgust from having to personally harm/kill an animal.

### Statistical analysis

All statistical analyses were performed using R version 4.3.3 (R Core Team, 2024). A Type II Wald chi-square ANOVAs were conducted using the “car” package (Fox & Weisberg, 2019), and post-hoc pairwise comparisons were conducted using the “emmeans” package (Lenth, 2025).

Generalised linear models (GLM) with a binomial family and a logit link were used to model the relationship between reports of bin-raiding foxes and participants’ urbanisation type, geographic region, and reported food provisioning activities. GLMs were also used to examine the likelihood of people reporting the use of fox control measures (professional services and DIY control methods) in relation to participant’s urbanisation type, geographic region, reported food provisioning activities, reported experience with bin-raiding foxes in their area (own property and/or neighbourhood), and their overall attitudes towards foxes. Finally, a GLM with a gaussian family and identity link was used to model participants’ overall attitudes towards foxes in relation to their urbanisation type, geographic region, reported food provisioning activities, and reported experience with bin-raiding foxes in their area. To assess the significance of individual predictors within each GLM, Type II Wald chi-square ANOVAs were conducted. For significant categorical predictors, post-hoc pairwise comparisons were performed using Tukey-adjusted p-values to explore differences between levels and the direction of effect within each significant factor.

## Results

### Participants’ urbanisation type, geographic region, and reports of human-fox interactions

Of the 1,373 people that responded to the survey, 9 people were excluded from the analyses because they were unable to correctly identify a fox when asked to click on the image of a fox from a lineup of different species. A further 89 people were excluded as their prior experience with bin-raiding foxes could not be determined due to missing responses. Information on urbanisation type, geographic region, and reports of human-fox interactions from the remaining 1,275 people are provided in Table 1 and Figure 1.

**Figure 1.**
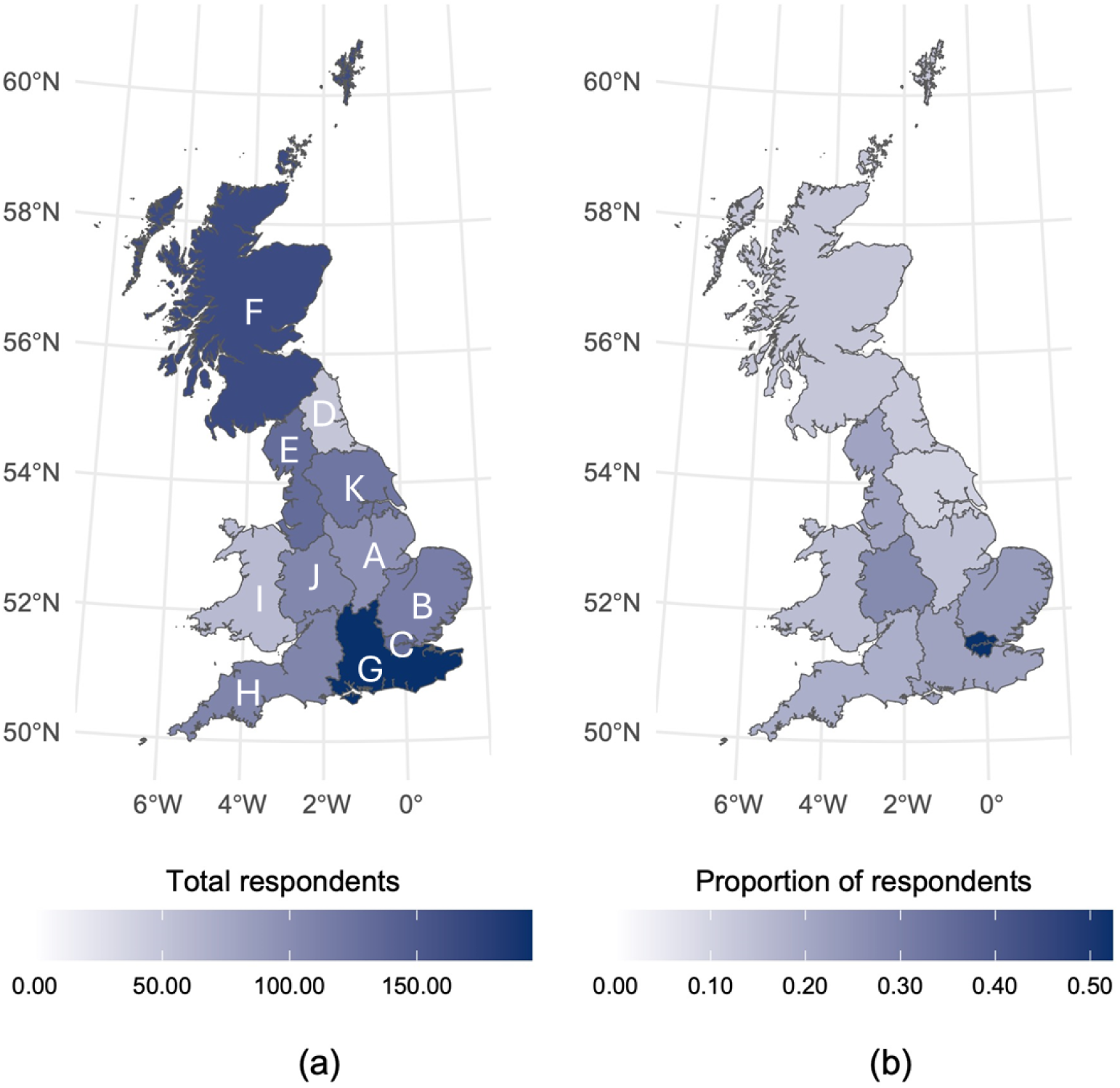
(a) The total number of respondents from each of the geographic regions within this study (N = 1,275 participants) and (b) the proportion of respondents that report bin-raiding behaviour, where ‘A’ is East Midlands (N = 97 participants), ‘B’ is East of England (N = 115 participants), ‘C’ is London (N = 130 participants), ‘D’ is North East (N = 49 participants), ‘E’ is North West (N = 131 participants), ‘F’ is Scotland (N = 164 participants), ‘G’ is South East (N = 195 participants), ‘H’ is South West (N = 108 participants), ‘I’ is Wales (N = 59 participants), ‘J’ is West Midlands (N = 105 participants) and ‘K’ is Yorkshire and the Humber (N = 122 participants).

**Table 1.**
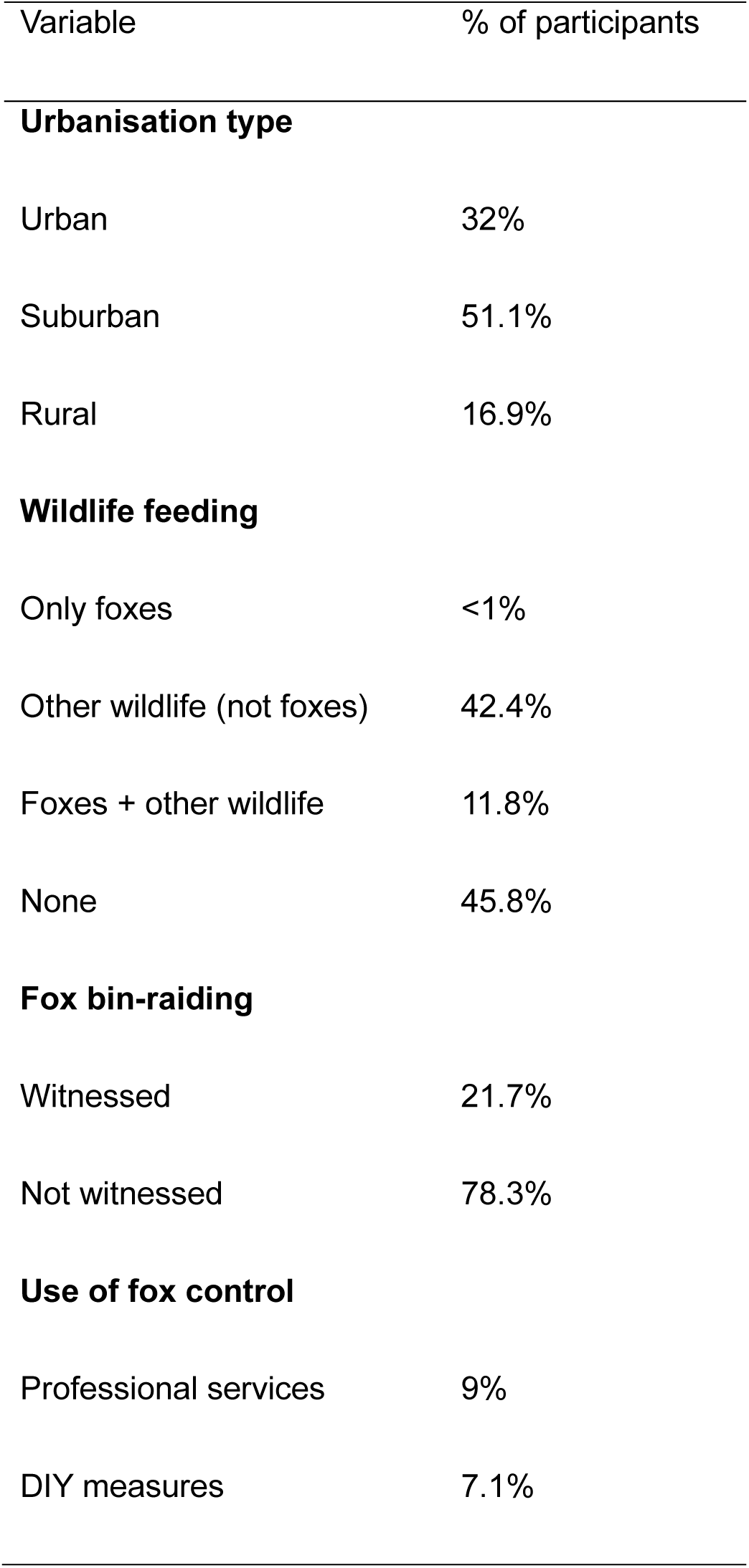
Information on participants’ reported urbanisation type, wildlife feeding habits, experience with fox bin-raiding, and use of fox control (N=1,275 participants).

| Variable | % of participants |
| --- | --- |
| **Urbanisation type** |  |
| Urban | 32% |
| Suburban | 51.1% |
| Rural | 16.9% |
| **Wildlife feeding** |  |
| Only foxes | <1% |
| Other wildlife (not foxes) | 42.4% |
| Foxes + other wildlife | 11.8% |
| None | 45.8% |
| **Fox bin-raiding** |  |
| Witnessed | 21.7% |
| Not witnessed | 78.3% |
| **Use of fox control** |  |
| Professional services | 9% |
| DIY measures | 7.1% |

### Spatial predictors of public wildlife feeding reports

Urbanisation type (χ²₂ = 21.277, p < 0.001), but not geographic region (χ²₁₀ = 7.558, p=0.672) were significantly related to reported wildlife feeding. Reports of people feeding wildlife were higher in urban compared to suburban (Z = −0.573, p < 0.001) and rural (Z = - 0.591, p < 0.001) areas, but suburban and rural areas did not differ (Z = −0.018, p =0.994).

### Predictors of fox bin-raiding reports

Urbanisation type (χ²₂ = 30.879, p < 0.001), public wildlife feeding (χ²₂ = 39.609, p < 0.001), and geographic region (χ²₁₀ = 49.707, p < 0.001) were significantly related to reported bin-raiding by foxes. People from urban areas were more likely to report bin-raiding foxes compared to people from suburban (Z = −4.004, p < 0.001) and rural (Z = −4.753, p < 0.001) areas (Figure 2). People who reported feeding foxes (either exclusively or along with other wildlife) were more likely to report bin-raiding than those who reported feeding other wildlife (not foxes) (Z = 6.307, p < 0.001) and no wildlife at all (Z = −4.913, p < 0.001) (Figure 3). People from London were significantly more likely to report bin-raiding foxes than those in eight of the other geographic regions (Table S1) (Figure 4).

**Figure 2.**
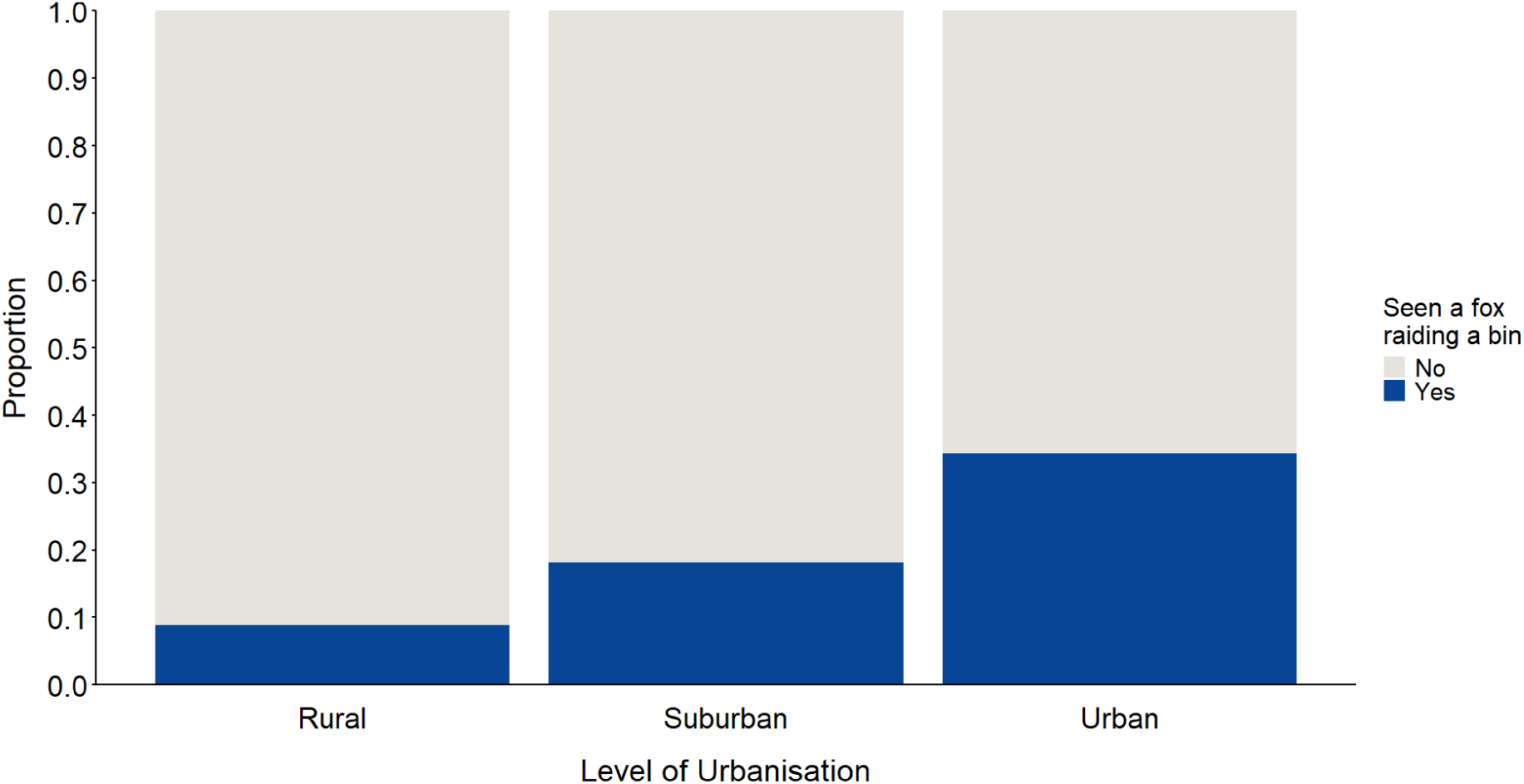
The proportion of people from each urbanisation type who reported seeing a fox raiding a bin, including rural (N = 216 participants), suburban (N = 651 participants) and urban (N = 408 participants) areas.

**Figure 3.**
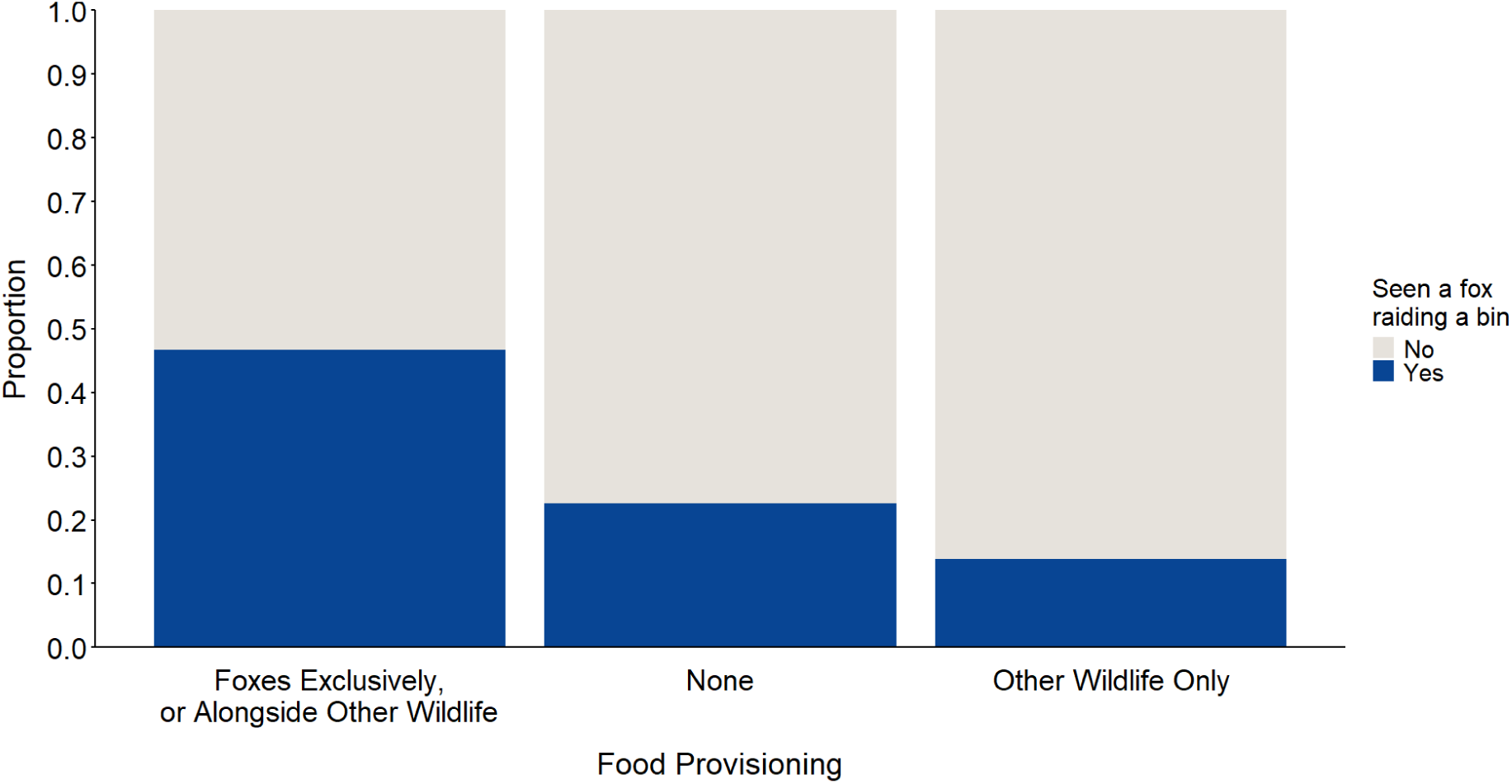
The proportion of people from each wildlife food provisioning category who reported seeing a fox raiding a bin, including people who reported feeding foxes exclusively or alongside other wildlife (N = 150 participants), people who reported feeding other wildlife (not foxes) (N = 541 participants), and people who did not report feeding any type of wildlife (N = 584 participants).

### Predictors of attitudes towards foxes

In terms of attitudes, 83.5% of participants were relatively more positive than negative towards foxes. Reports of food provisioning (χ²₂ = 41.095, p < 0.001) and fox bin-raiding experience (χ²_1_ = 9.208, p = 0.002) were significantly related to public attitudes, whereas urbanisation type (χ²₂ = 0.515, p = 0.773) and geographic region (χ²_10_ = 16.129, p = 0.096) were not. People who reported feeding foxes (either alone or with other wildlife) were more likely to have more positive attitudes towards foxes than people who did not report feeding any wildlife (Z = −2.419, p = 0.042); however, their attitudes did not significantly differ from people who reported feeding other wildlife (not foxes) (Z = -1.683, p = 0.212). People who reported witnessing fox bin-raiding had poorer attitudes than people who had not (Z = 3.035, p = 0.003).

### Predictors of fox control

#### Participants’ reported use of professional pest control services

People’s reported use of professional services to resolve a fox-related issue within their area was significantly associated with urbanisation type (χ²₂ 6.019, p = 0.049), degree of reported food provisioning (χ²₂ = 29.456, p < 0.001), reported fox bin-raiding experience (χ²₁ = 18.059, p < 0.001) and attitudes towards foxes (χ²₁ = 11.182, p < 0.001). Geographic region was not significantly related to the reported use of professional services (χ²_10_ = 9.016, p = 0.531).

People from urban areas were more likely to report having contacted professional services than those from suburban areas (Z = −2.384, p = 0.045); however, comparisons between rural and suburban (Z = 1.238, p = 0.430), and rural and urban areas (Z = −0.497, p = 0.873) were not significant. People who reported feeding foxes (either alone or with other wildlife) were more likely to contact professional services than those who reported feeding other wildlife (not foxes) (Z = 4.720, p < 0.001) and no wildlife at all (Z = -5.219, p < 0.001; Figure S1). People who reported having previously witnessed bin-raiding by foxes were more likely to contact professional services than people who had not (Z = −4.311, p < 0.001; Figure S2). Finally, people with more negative attitudes were more likely to report having contacted professional services compared to people with more positive attitudes (Z = -3.373, p < 0.001).

#### Participants’ reported use of DIY measures to control foxes

People’s reported use of DIY measures to resolve a fox-related issue within their area was significantly associated with their degree of reported food provisioning (χ²₂ = 30.360, p < 0.001), geographic region (χ²₁₀ = 19.332, p = 0.036), reported fox bin-raiding experience (χ²₁ = 12.752, p < 0.001), and attitudes towards foxes (χ²₁ = 13.093, p < 0.001). Urbanisation type was not significantly associated with the reported use of DIY control measures (χ²₂ = 5.018, p = 0.081).

People who reported feeding foxes (either alone or with other wildlife) were more likely to use DIY control measures than those who reported feeding other wildlife (not foxes) (Z = 5.255, p < 0.001) and no wildlife at all (Z = −4.534, p < 0.001) (Figure S3 and S4). People who reported having previously witnessed bin-raiding by foxes were more likely to use DIY control measures than people who had not (Z = -3.621, p < 0.001). People with more negative attitudes were more likely to report using DIY control measures compared to people with more positive attitudes (Z = -3.639, p < 0.001). Finally, none of the pair-wise comparisons between different geographic regions of the UK were significant (Table S2).

## Discussion

The current study examined how urbanisation type and geographic region are related to people’s reports of bin-raiding, food provisioning habits, attitudes, and reported use of control measures towards wild red foxes. Reports of bin-raiding by foxes were associated with higher reported levels of public wildlife feeding, more negative attitudes towards foxes, and greater reported use of professional and DIY fox control measures. However, the role of urbanisation type and geographic differences varied among these relationships, highlighting the complex interaction between spatial factors and the subjective human dimensions of wildlife interactions, including public attitudes and self-reported encounters with foxes.

Previous studies have found links between feeding wildlife and people’s tolerance of those species (Baker et al., 2020; Puri et al., 2024). Consistent with these other studies, the current study found that people who reported feeding wildlife were also more likely to have positive attitudes towards foxes. Although we are unable to attribute causality within the current study, feeding wildlife, including foxes, may encourage people to have a closer connection to nature (Cox & Gaston, 2018; White et al., 2023). Alternatively, people with more positive attitudes may be more likely to feed foxes and other wildlife. We encourage future research to test these and other possibilities, ideally using experimentally controlled field tests.

Although people who reported feeding wildlife were more tolerant of foxes, the current study also found that such practices were related to a greater likelihood of reporting bin-raiding behaviour and using fox control measures. Without controlled experimental tests, it is unclear whether public wildlife feeding encourages bin-raiding behaviour in foxes, or whether the visibility of bin-raiding foxes encourages people to interact with them by feeding them (Baker et al., 2004). Moreover, our findings could potentially reflect a complex relationship between feeding behaviour and attitudes, where people who feed foxes have more positive views of them, but feeding may also lead to increase contact and conflict (Cahill et al., 2012; Cox & Gaston, 2018; Kumar et al., 2019; Schell et al., 2021), prompting some to use control measures. Finally, it is also possible that feeding serves different functions for different individuals – from expressing affection for wildlife to baiting them for pest control (Cox & Gaston, 2016; Taggart et al., 2023). Thus, people who report feeding foxes may not be a homogenous group and could, instead, represent a myriad of motivations for interacting with foxes. Further research is needed to test these and other possibilities. Further work should also investigate whether respondents who do *not* feed wildlife observe fox bin-raiding and engage in pest control because their neighbours are feeding them.

Bin-raiding foxes were reported more frequently in urban areas compared to suburban and rural areas, particularly the UK’s largest city, London. Such findings may reflect the higher likelihood of encounters and ease of observation of foxes in densely populated urban areas (Morton et al. 2023; Scott et al., 2014). Alternatively, such findings may reflect the fact that urban foxes are likely exposed to more rubbish, more bins, and are typically bolder and more willing to physically explore human-made objects containing food compared to rural populations (Morton et al., 2023). This latter interpretation would also be consistent with a growing body of research highlighting significant behavioural differences, including greater boldness and exploitation of anthropogenic food sources, among urban populations compared to rural conspecifics in a wide variety of taxa, including other carnivores (Breck et al., 2019; Griffin et al., 2017; Murray et al., 2015). Finally, further research is needed to assess whether people’s perceptions of foxes accurately reflect fox behaviour in real life. For example, comparing subjective household questionnaires with objective ecological data (e.g., trail camera footage of foxes) could help rule out perceived (not real) fox-related events arising from mistaken identity or attribution due to signs left by other bin-raiding animals (e.g., cats or birds).

Studies show that people with more positive attitudes and experiences with a species are generally more willing to co-exist with them (Baker et al., 2020; Hosaka et al., 2017; Mohamad Muslim et al., 2018; Puri et al., 2024; Sweet et al., 2024). Consistent with this notion, the current study found that reports of fox control measures were more common among people with more negative attitudes towards foxes and among those who reported having observed bin-raiding in their area. These findings also provide support to the notion that negative experiences with wildlife can impact public tolerance (Baker et al., 2020; Conejero et al., 2019; Keziah J Hobson et al., 2024; Naughton-Treves et al., 2003; Puri et al., 2024). As discussed, experimentally controlled tests are needed to established causality among the associations identified in this study.

Spatial factors contributed to people’s likelihood of reporting the use of professional and DIY fox control measures. Specifically, people living in suburban areas were less likely to report using professional services than those in urban environments, aligning with higher reported rates of bin-raiding in urban locations. Human-nature connectedness tends to be weaker among urban residents compared to other people (Barragan-Jason et al., 2025; Kageyama et al., 2024; Morton & Soulsbury, 2025; Richardson et al., 2022), which may lead urban residents to perceive negative interactions, such as fox bin-raiding, as a more pressing issue in need of control or professional advice. Moreover, people in urban areas have also been found to believe that foxes cause more problems in general (Baker et al., 2020), which could further explain the higher reported use of professional services. However, in contrast, urbanisation type did not significantly predict reports of DIY control, suggesting that this form of response may be related to other contextual or individual factors.

Geographic region was not significantly associated with people’s likelihood of reporting the use of professional services to resolve a fox-related issue within their area, indicating that regional variation may play a lesser role than urbanisation in shaping this particular behaviour.

This aligns with Baker et al. (2020), who found no association between geographic region and past use of control measures (both professional and DIY) for several ‘nuisance’ species. However, in the current study, geographical region was a significant overall predictor of DIY control, though no individual pairwise comparisons reached significance. These results may reflect the lower statistical power of multiple comparisons, but could also indicate broad geographic differences that are not fully captured by administrative boundaries. Collectively, these findings suggest that spatial context predicts variation in the use of control measures, but the strength and nature of these associations may vary depending on the type of action taken. Future research would benefit from exploring alternative spatial configurations, such as ecological or cultural boundaries, which may capture these dynamics better. Finally, although professional services are widespread throughout the UK, further research should investigate whether geographic variation in the reported use of DIY measures is explained by factors limiting the use or accessibility of professional services, including cultural or socio-economic factors.

Spatial factors predicted reports of bin-raiding, wildlife food provisioning, and use of different fox control measures, but they did not predict public attitudes, suggesting a spatial uncoupling between attitudes and people’s reported interactions with foxes. One possibility is that public attitudes may be shaped by broader societal narratives about foxes, or social norms shared among UK residents about wildlife more generally (Dickman, 2010). However, the fact that attitudes were still linked to participants’ observations of bin-raiding and the use of fox control measures implies that personal experiences could still reinforce or modify these broader perceptions. Further research is needed to better understand how personal experiences and popular culture biases might interact to shape public attitudes towards foxes, which has been found in a number of other species (Bombieri et al., 2018; Nanni et al., 2020; Sabatier & Huveneers, 2018). We also recommend that future research extend our findings by including a larger sample of participants using pest control measures. Finally, we recommend integrating additional measures of urbanisation and other landscape features beyond the three-level classification (urban, suburban, rural) used here.

### Implications for urban rewilding

Although further research is needed, our findings potentially offer an example of why urban rewilding initiatives may benefit from obtaining local spatial insights from subjective human dimensions, rather than relying solely on broad, generalised trends across communities. For instance, as discussed, most pro-environmental messages related to human-wildlife coexistence are broadly communicated across communities without accounting for spatial variation in people’s subjective experiences with those species. This is noteworthy because, as discussed, socio-psychological research shows that people are unlikely to change their behaviour unless it aligns with what they already know, value, or believe (Toomey, 2023). In the case with foxes, our study and others suggest that these animals are not the ‘prolific bin-raiders’ commonly portrayed by popular culture (Morton et al., 2023) and so promoting this type of message to a community that already believes or reports having such experiences may not be persuasive and could, instead, undermine trust or lead to more entrenched negative views of that species (Toomey, 2023). By understanding local human dimensions and incorporating them into communication strategies, rewilding programmes can potentially help to avoid issues that may arise from circulating generalised pro-environmental messages that do not align with their specific local community’s attitudes or perceived experiences.

## Conclusion

This study highlights the complex interaction between spatial factors and the subjective human dimensions of human-wildlife interactions. Specifically, we demonstrate that urbanisation and geographic region can predict variation in the likelihood of people reporting wildlife encounters and the use of control measures. Thus, without local context, generalised assumptions are potentially insufficient for predicting how communities may interact and behave towards a species, which could undermine one-size-fits-all strategies to encourage behavioural change in people to promote coexistence. Because of this complexity, our study underscores the value of adopting locally informed approaches to conservation, including urban rewilding initiatives, to account for possible localised spatial variation in these subjective human dimensions.

## Data Availability

All data are provided in the supplementary materials (Dataset S1).

## Acknowledgments

We thank the participants in this study, and everyone involved with the *British Carnivore Project*. K.A.A. was supported at the time of writing by a scholarship from the

Doctoral College at the University of Hull. F.B.M. thanks the University of Hull and the UKRI Natural Environment Research Council (NERC) (Grant No. NE/X018342/1) for funding.

## References

1. Aronson, M. F., La Sorte, F. A., Nilon, C. H., Katti, M., Goddard, M. A., Lepczyk, C. A., Warren, P. S., Williams, N. S., Cilliers, S., & Clarkson, B. (2014). A global analysis of the impacts of urbanization on bird and plant diversity reveals key anthropogenic drivers. Proceedings of the Royal Society B: Biological Sciences, 281(1780), 20133330.

2. Arup, & C40 Cities. (2022). Urban Rewilding The value and co-benefits of nature in urban spaces. https://www.arup.com/insights/urban-rewilding-the-value-and-co-benefits-ofnature-in-urban-spaces/

3. Baker, Maw, S. A., Johnson, P. J., & Macdonald, D. W. (2020). Not in my backyard: Public perceptions of wildlife and ’pest control’ in and around UK homes, and local authority ’pest control’ Animals, 10, 222.

4. Baker, P., Funk, S., Harris, S., Newman, T., Saunders, G., & White, P. (2004). The impact of human attitudes on the social and spatial organization of urban foxes (Vulpes vulpes) before and after an outbreak of sarcoptic mange. Proceedings 4th International Urban Wildlife Symposium 153–163.

5. Baker, P., Harris, S., & White, P. (2006). After the hunt: the future for foxes in Britain. International Fund for Animal Welfare.

6. Balčiauskas, L., & Kazlauskas, M. (2014). Forty years after reintroduction in a suboptimal landscape: public attitudes towards European bison. European Journal of Wildlife Research, 60(1), 155–158.

7. Barrable, A., & Booth, D. (2022). Disconnected: what can we learn from individuals with very low nature connection? International journal of environmental research and public health, 19(13), 8021.

8. Barragan-Jason, G., Cauchoix, M., Diaz-Valencia, P. A., Syssau-Vaccarella, A., Hemet, S., Cardozo, C., Skevington, S. M., Heeb, P., & Parmesan, C. (2025). Human–nature connectedness and sustainability across lifetimes: A comparative cross-sectional study in France and Colombia. People and Nature, 7(1), 99–111.

9. Barrows, P. D., Richardson, M., Hamlin, I., & Van Gordon, W. (2022). Nature connectedness, nonattachment, and engagement with nature’s beauty predict pro-nature conservation behavior. Ecopsychology, 14(2), 83–91.

10. Basak, S. M., Hossain, M. S., O’Mahony, D. T., Okarma, H., Widera, E., & Wierzbowska, I. A. (2022). Public perceptions and attitudes toward urban wildlife encounters–A decade of change. Science of the Total Environment, 834, 155603.

11. Bijl, H., & Csányi, S. (2022). The reasons for the range expansion of the grey wolf, coyote and red fox. Review on Agriculture and Rural Development, 11(1-2), 46–53.

12. Bjerke, T., Ødegårdstuen, T. S., & Kaltenborn, B. P. (1998). Attitudes toward animals among Norwegian children and adolescents: Species preferences. Anthrozoos, 11(4), 227235.

13. Bombieri, G., Nanni, V., Delgado, M. d. M., Fedriani, J. M., López-Bao, J. V., Pedrini, P., & Penteriani, V. (2018). Content analysis of media reports on predator attacks on humans: Toward an understanding of human risk perception and predator acceptance. BioScience, 68(8), 577–584.

14. Brand, A., & Baldwin, M. (2020). Public attitudes to urban foxes in London and the south east. Mammal Society.

15. Breck, S. W., Poessel, S. A., Mahoney, P., & Young, J. K. (2019). The intrepid urban coyote: a comparison of bold and exploratory behavior in coyotes from urban and rural environments. Scientific Reports, 9, 2104.

16. Bridge, B., & Harris, S. (2020). Do urban red foxes attack people? An exploratory study and review of incidents in Britain. Human-Wildlife Interactions, 14(2), 151–165.

17. Bruskotter, J. T., & Fulton, D. C. (2012). Will hunters steward wolves? A comment on Treves and Martin. Society & Natural Resources, 25(1), 97–102.

18. Bruskotter, J. T., Singh, A., Fulton, D. C., & Slagle, K. (2015). Assessing tolerance for wildlife: clarifying relations between concepts and measures. Human Dimensions of Wildlife, 20(3), 255–270.

19. Bruskotter, J. T., & Wilson, R. S. (2014). Determining where the wild things will be: Using psychological theory to find tolerance for large carnivores Conservation Letters, 7, 158165.

20. Cahill, S., Llimona, F., Cabañeros, L., & Calomardo, F. (2012). Characteristics of wild boar (*Sus scrofa*) habituation to urban areas in the Collserola Natural Park (Barcelona) and comparison with other locations. Animal Biodiversity and Conservation, 35(2), 221233.

21. Carver, S., Convery, I., Hawkins, S., Beyers, R., Eagle, A., Kun, Z., Van Maanen, E., Cao, Y., Fisher, M., & Edwards, S. R. (2021). Guiding principles for rewilding. Conservation Biology, 35(6), 1882–1893.

22. Castañeda, I., Doherty, T. S., Fleming, P. A., Stobo-Wilson, A. M., Woinarski, J. C., & Newsome, T. M. (2022). Variation in red fox *Vulpes vulpes* diet in five continents. Mammal Review, 52(3), 328–342.

23. Cazalis, V., Loreau, M., & Barragan-Jason, G. (2023). A global synthesis of trends in human experience of nature. Frontiers in Ecology and the Environment, 21(2), 85–93.

24. Conejero, C., Castillo-Contreras, R., González-Crespo, C., Serrano, E., Mentaberre, G., Lavín, S., & López-Olvera, J. R. (2019). Past experiences drive citizen perception of wild boar in urban areas. Mammalian Biology, 96, 68–72.

25. Contesse, P., Hegglin, D., Gloor, S., Bontadina, F., & Deplazes, P. (2004). The diet of urban foxes (*Vulpes vulpes*) and the availability of anthropogenic food in the city of Zurich, Switzerland. Mammalian biology, 69(2), 81–95.

26. Cox, D. T., & Gaston, K. J. (2016). Urban bird feeding: Connecting people with nature. PLoS ONE, 11(7), e0158717.

27. Cox, D. T., & Gaston, K. J. (2018). Human–nature interactions and the consequences and drivers of provisioning wildlife. Philosophical Transactions of the Royal Society B: Biological Sciences, 373(1745), 20170092.

28. Cox, D. T., Hudson, H. L., Shanahan, D. F., Fuller, R. A., & Gaston, K. J. (2017). The rarity of direct experiences of nature in an urban population. Landscape and Urban Planning, 160, 79–84.

29. Cox, D. T., Shanahan, D. F., Hudson, H. L., Plummer, K. E., Siriwardena, G. M., Fuller, R. A., Anderson, K., Hancock, S., & Gaston, K. J. (2017). Doses of neighborhood nature: the benefits for mental health of living with nature. AIBS Bulletin, 67(2), 147–155.

30. Dickman, A. J. (2010). Complexities of conflict: the importance of considering social factors for effectively resolving human–wildlife conflict. Animal Conservation, 13(5), 458–466.

31. Dunn, M., Marzano, M., Forster, J., & Gill, R. M. (2018). Public attitudes towards “pest” management: Perceptions on squirrel management strategies in the UK. Biological Conservation, 222, 52–63.

32. Finnerty, P. B., Carthey, A. J., Banks, P. B., Brewster, R., Grueber, C. E., Houston, D., Martin, J. M., McManus, P., Roncolato, F., & van Eeden, L. M. (2025). Urban rewilding to combat global biodiversity decline. BioScience, biaf062.

33. Fox, J., & Weisberg, S. (2019). An R Companion to Applied Regression*, Third edition. Sage,* Thousand Oaks CA. In https://www.john-fox.ca/Companion/

34. Gaston, K. J., & Soga, M. (2020). Extinction of experience: The need to be more specific. People and Nature, 2(3), 575–581.

35. Gianotti, A. G. S., Getson, J. M., Hutyra, L. R., & Kittredge, D. B. (2016). Defining urban, suburban, and rural: a method to link perceptual definitions with geospatial measures of urbanization in central and easter Massachusetts. Urban Ecosystems, 19, 823–833.

36. Griffin, A. S., Netto, K., & Peneaux, C. (2017). Neophilia, innovation and learning in an urbanised world: a critical evaluation of mixed findings. Current Opinion in Behavoral Sciences, 16, 15–22.

37. Griffiths-Lee, J., Nicholls, E., & Goulson, D. (2022). Sown mini-meadows increase pollinator diversity in gardens. Journal of Insect Conservation, 26(2), 299–314.

38. Harris. (1981). The food of suburban foxes (Vulpes vulpes), with special reference to London. Mammal Review, 11, 151–168.

39. Hill, M. J., Biggs, J., Thornhill, I., Briers, R. A., Gledhill, D. G., White, J. C., Wood, P. J., & Hassall, C. (2017). Urban ponds as an aquatic biodiversity resource in modified landscapes. Global Change Biology, 23(3), 986–999.

40. Hobson, K. J., Stringer, A., Gill, R., MacPhearson, J., & Lambin, X. (2024). Interests, beliefs, experience and perceptions shape tolerance towards impacts of recovering predators People and Nature, 6, 117–133.

41. Hobson, K. J., Stringer, A., Gill, R., MacPherson, J., & Lambin, X. (2024). Interests, beliefs, experience and perceptions shape tolerance towards impacts of recovering predators. People and Nature, 6(1), 117–133.

42. Hosaka, T., Sugimoto, K., & Numata, S. (2017). Childhood experience of nature influences the willingness to coexist with biodiversity in cities. Palgrave Communications, 3(1), 18.

43. Kageyama, S., Saito, T., Tajima, Y., & Hashimoto, S. (2024). Human–nature connectedness is positively correlated with the perceived value of nature regardless of urbanization levels. Sustainability Science, 1–18.

44. Kansky, R., Kidd, M., & Knight, A. T. (2014). Meta-analysis of attitudes toward damage-causing mammalian wildlife. Conservation Biology, 28(4), 924–938.

45. Kimmig, S. E., Flemming, D., Kimmerle, J., Cress, U., & Brandt, M. (2020). Elucidating the socio-demographics of wildlife tolerance using the example of the red fox (*Vulpes vulpes*) in Germany. Conservation Science and Practice, 2(7), e212.

46. König, H. J., Kiffner, C., Kramer-Schadt, S., Fürst, C., Keuling, O., & Ford, A. T. (2020). Human–wildlife coexistence in a changing world. Conservation Biology, 34(4), 786794.

47. Kumar, N., Jhala, Y. V., Qureshi, Q., Gosler, A. G., & Sergio, F. (2019). Human-attacks by an urban raptor are tied to human subsidies and religious practices. Scientific Reports, 9(1), 2545.

48. Lengieza, M. L., Aviste, R., & Richardson, M. (2023). The human–nature relationship as a tangible target for pro-environmental behaviour—Guidance from interpersonal relationships. Sustainability, 15(16), 12175.

49. Lengieza, M. L., Richardson, M., & Hughes, J. P. (2025). Feature networks: The environmental features that are central to nature-connectedness experiences. Landscape and Urban Planning, 259, 105362.

50. Lenth, R. (2025). emmeans: Estimated Marginal Means, aka Least-Squares Means*. R package version 1.10.7*. In https://CRAN.R-project.org/package=emmeans

51. Li, G., Fang, C., Li, Y., Wang, Z., Sun, S., He, S., Qi, W., Bao, C., Ma, H., & Fan, Y. (2022). Global impacts of future urban expansion on terrestrial vertebrate diversity. Nature Communications, 13(1), 1628.

52. Marshall, C. A., Wilkinson, M. T., Hadfield, P. M., Rogers, S. M., Shanklin, J. D., Eversham, B. C., Healey, R., Kranse, O. P., Preston, C. D., & Coghill, S. J. (2023). Urban wildflower meadow planting for biodiversity, climate and society: An evaluation at King’s College, Cambridge. Ecological Solutions and Evidence, 4(2), e12243.

53. Martyn, P., & Brymer, E. (2016). The relationship between nature relatedness and anxiety. Journal of health psychology, 21(7), 1436–1445.

54. McCleery, R. A., Moorman, C. E., Wallace, M. C., & Drake, D. (2012). Managing Urban Environments for Wildlife. In N. J. Silvy (Ed.), The Wildlife Techniques Manual: Management (Vol. 2, pp. 169–191). Johns Hopkins University Press.

55. McEwan, K., Richardson, M., Sheffield, D., Ferguson, F. J., & Brindley, P. (2019). A smartphone app for improving mental health through connecting with urban nature. International journal of environmental research and public health, 16(18), 3373.

56. McKinney, M. L. (2002). Urbanization, biodiversity, and conservation: the impacts of urbanization on native species are poorly studied, but educating a highly urbanized human population about these impacts can greatly improve species conservation in all ecosystems. BioScience, 52(10), 883–890.

57. McKinney, M. L. (2006). Urbanization as a major cause of biotic homogenization. Biological Conservation, 127(3), 247–260.

58. McPherson, S. C., Sumasgutner, P., Hoffman, B. H., Padbury, B. D., Brown, M., Caine, T. P., & Downs, C. T. (2021). Surviving the urban jungle: Anthropogenic threats, wildlife-conflicts, and management recommendations for African crowned eagles. Frontiers in Ecology and Evolution, 9, 662623.

59. Moesch, S. S., Jeschke, J. M., Lokatis, S., Peerenboom, G., Kramer-Schadt, S., Straka, T. M., & Haase, D. (2024). The frequent five: Insights from interviews with urban wildlife professionals in Germany. People and Nature, 6(5), 2091–2108.

60. Moesch, S. S., Straka, T. M., Jeschke, J. M., Haase, D., & Kramer-Schadt, S. (2024). The good, the bad, and the unseen: wild mammal encounters influence wildlife preferences of residents across socio-demographic gradients. Ecology and Society, 29.

61. Mohamad Muslim, H. F., Tetsuro, H., Shinya, N., & Yahya, N. A. (2018). Nature experience promotes preference for and willingness to coexist with wild animals among urban and suburban residents in Malaysia. Ecological Processes, 7, 1–12.

62. Morton, F. B., Gartner, M., Norrie, E.-M., Haddou, Y., Soulsbury, C. D., & Adaway, K. A. (2023). Urban foxes are bolder but not more innovative than their rural conspecifics Animal Behaviour, 203, 101–113.

63. Morton, F. B., Gartner, M., Norrie, E.-M., Haddou, Y., Soulsbury, C. D., & Adaway, K. A. (2023). Urban foxes are bolder but not more innovative than their rural conspecifics. Animal Behaviour, 203, 101–113.

64. Morton, F. B., Henri, D., Adaway, K. A., Soulsbury, C. D., & Hopkins, C. R. (2024). Communicating information about the psychology of a wild carnivore, the red fox, influences perceived attitudinal changes but not overall tolerance in people. Biological Conservation, 296, 110653.

65. Morton, F. B., & Soulsbury, C. D. (2025). Experiencing the wild: red fox encounters are related to stronger nature connectedness, not anxiety, in people. Human Dimensions of Wildlife, 1–18.

66. Murray, M., Cembrowski, A., Latham, A. D. M., Lukasik, V. M., Pruss, S., & St Clair, C. C. (2015). Greater consumption of protein-poor anthropogenic food by urban relative to rural coyotes increases diet breadth and potential for human–wildlife conflict. Ecography, 38, 1235–1242.

67. Nanni, V., Caprio, E., Bombieri, G., Schiaparelli, S., Chiorri, C., Mammola, S., Pedrini, P., & Penteriani, V. (2020). Social media and large carnivores: Sharing biased news on attacks on humans. Frontiers in Ecology and Evolution, 8, 71.

68. Naughton-Treves, L., Grossberg, R., & Treves, A. (2003). Paying for tolerance: rural citizens’ attitudes toward wolf depredation and compensation. Conservation Biology, 17(6), 1500–1511.

69. Nyhus, P. J. (2016). Human–wildlife conflict and coexistence. Annual Review of Environment and Resources, 41(1), 143–171.

70. Ostermann-Miyashita, E.-F., Pernat, N., Koenig, H. J., Hemminger, K., Gandl, N., BellingrathKimura, S. D., Hibler, S., & Kiffner, C. (2023). Attitudes of wildlife park visitors towards returning wildlife species: An analysis of patterns and correlates. Biological Conservation, 278, 109878.

71. Otto, S., & Pensini, P. (2017). Nature-based environmental education of children: Environmental knowledge and connectedness to nature, together, are related to ecological behaviour. Global environmental change, 47, 88–94.

72. Pettorelli, N., Dancer, A. D., Durant, S. M., Hoffmann, M., Laughlin, B., Pilkington, J., Pecorelli, J., Seiffert, S., Shadbolt, T., & Terry, A. (2022). Rewilding our cities.

73. Puri, M., Johannsen, K. L., Goode, K. O., & Pienaar, E. F. (2024). Addressing the challenge of wildlife conservation in urban landscapes by increasing human tolerance for wildlife. People and Nature, 6(3), 1116–1129.

74. R Core Team. (2024). R: A Language and Environment for Statistical Computing. R Foundation for Statistical Computing. In https://www.R-project.org/

75. Ramadhan, A. L. (2024). Understanding Human-Wildlife Interactions in Urban Environments: Implications for Conflicts, Disease Transmission, and Conservation. Law and Economics, 18(2), 99–109.

76. Richardson, M., Hamlin, I., Elliott, L. R., & White, M. P. (2022). Country-level factors in a failing relationship with nature: Nature connectedness as a key metric for a sustainable future. Ambio, 51(11), 2201–2213.

77. Richardson, M., Passmore, H. A., Barbett, L., Lumber, R., Thomas, R., & Hunt, A. (2020). The green care code: How nature connectedness and simple activities help explain pro-nature conservation behaviours. People and Nature, 2(3), 821–839.

78. Sabatier, E., & Huveneers, C. (2018). Changes in media portrayal of human-wildlife conflict during successive fatal shark bites. Conservation and Society, 16(3), 338–350.

79. Sato, Y. (2017). The future of urban brown bear management in Sapporo, Hokkaido, Japan: a review. Mammal Study, 42(1), 17–30.

80. Schell, C. J., Stanton, L. A., Young, J. K., Angeloni, L. M., Lambert, J. E., Breck, S. W., & Murray, M. H. (2021). The evolutionary consequences of human–wildlife conflict in cities. Evolutionary Applications, 14(1), 178–197. 10.1111/eva.13131

81. Scott, D. M., Berg, M. J., Tolhurst, B. A., Chauvenet, A. L., Smith, G. C., Neaves, K., Lochhead, J., & Baker, P. J. (2014). Changes in the distribution of red foxes (*Vulpes vulpes*) in urban areas in Great Britain: findings and limitations of a media-driven nationwide survey. PLoS ONE, 9(6), e99059.

82. Sibthorpe, R. L., & Brymer, E. (2020). Disconnected from nature: the lived experience of those disconnected from the natural world. Innovations in a changing world, 59.

83. Sidemo-Holm, W., Ekroos, J., Reina García, S., Söderström, B., & Hedblom, M. (2022). Urbanization causes biotic homogenization of woodland bird communities at multiple spatial scales. Global Change Biology, 28(21), 6152–6164.

84. Simkin, R. D., Seto, K. C., McDonald, R. I., & Jetz, W. (2022). Biodiversity impacts and conservation implications of urban land expansion projected to 2050. Proceedings of the National Academy of Sciences, 119(12), e2117297119.

85. Soe, E., Davison, J., Süld, K., Valdmann, H., Laurimaa, L., & Saarma, U. (2017). Europe-wide biogeographical patterns in the diet of an ecologically and epidemiologically important mesopredator, the red fox *Vulpes vulpes*: a quantitative review. Mammal Review, 47(3), 198–211.

86. Soga, M., & Gaston, K. J. (2016). Extinction of experience: the loss of human–nature interactions. Frontiers in Ecology and the Environment, 14(2), 94–101.

87. Sol, D., González-Lagos, C., Moreira, D., Maspons, J., & Lapiedra, O. (2014). Urbanisation tolerance and the loss of avian diversity. Ecology letters, 17(8), 942–950.

88. Soulsbury, C. D., Baker, P. J., Iossa, G., & Harris, S. (2010). Red Foxes *Vulpes vulpes*. In S. D. Gehrt, S. P. Riley, & B. L. Cypher (Eds.), Urban carnivores (pp. 63–78). Johns Hopkins university press.

89. Soulsbury, C. D., Iossa, G., Baker, P. J., Cole, N. C., Funk, S. M., & Harris, S. (2007). The impact of sarcoptic mange Sarcoptes scabiei on the British fox Vulpes vulpes population. Mammal Review, 37(4), 278–296. 10.1111/j.13652907.2007.00101.x

90. Soulsbury, C. D., & White, P. C. (2015). Human-wildlife interactions in urban areas: A review of conflicts, benefits and opportunities. Wildlife Research, 42, 541–553.

91. Straka, T. M., Glahe, C., Dietrich, U., Bui, M., & Kowarik, I. (2025). From nature experience to pro-conservation action: How generational amnesia and declining nature-relatedness shape behaviour intentions of adolescents and adults. Ambio, 1–20.

92. Sweet, F. S., Mimet, A., Shumon, M. N. U., Schirra, L. P., Schäffler, J., Haubitz, S. C., Noack, P., Hauck, T. E., & Weisser, W. W. (2024). There is a place for every animal, but not in my back yard: a survey on attitudes towards urban animals and where people want them to live. Journal of Urban Ecology, 10(1), juae006.

93. Taggart, P. L., Taylor, P., Patel, K. K., & Noble, D. W. (2023). Baiting in conservation and pest management: a systematic review of its global applications in a changing world. Biological Conservation, 284, 110214.

94. Tan, A. S., de la Torre, J. A., Wong, E. P., Thuppil, V., & Campos-Arceiz, A. (2020). Factors affecting urban and rural tolerance towards conflict-prone endangered megafauna in Peninsular Malaysia. Global Ecology and Conservation, 23, e01179.

95. The_Humane_Society_of_the_United_States. (2023). Conflict resolution guide: Humane wildlife conflict resolution guide. HumanePro. https://humanepro.org/sites/default/files/documents/HSUS_HumaneWildlifeConflictResolutionGuide_2023.pdf

96. Threlfall, C., Mata, L., Mackie, J., Hahs, A. K., Stork, N. E., Williams, N. S., & Livesley, S. J. (2017). Increasing biodiversity in designed urban green spaces through simple vegetation interventions. 2017 ESA Annual Meeting (August 6--11),

97. Toomey, A. H. (2023). Why facts don’t change minds: Insights from cognitive science for the improved communication of conservation research. Biological Conservation, 278, 109886.

98. United Nations, D. o. E. a. S. A., Population Division. (2019). World Urbanization Prospects 2018: Highlights (ST/ESA/SER. A/421).

99. Vantassel, S. M., & Groepper, S. R. (2015). A survey of wildlife damage management techniques used by wildlife control operators in urbanized environments in the USA. In F. Angelici (Ed.), Problematic wildlife: A cross-disciplinary approach (pp. 175–204).

100. Webb, J., & Moxon, S. (2023). A study protocol to understand urban rewilding behaviour in relation to adaptations to private gardens. Cities & health, 7(2), 273–281.

101. Whitburn, J., Linklater, W., & Abrahamse, W. (2020). Meta-analysis of human connection to nature and proenvironmental behavior. Conservation Biology, 34(1), 180–193.

102. White, M. E., Hamlin, I., Butler, C. W., & Richardson, M. (2023). The joy of birds: The effect of rating for joy or counting garden bird species on wellbeing, anxiety, and nature connection. Urban Ecosystems, 26(3), 755–765.

103. White, M. P., Elliott, L. R., Grellier, J., Economou, T., Bell, S., Bratman, G. N., Cirach, M., Gascon, M., Lima, M. L., & Lõhmus, M. (2021). Associations between green/blue spaces and mental health across 18 countries. Scientific Reports, 11(1), 8903.

104. Wooster, E., Fleck, R., Torpy, F., Ramp, D., & Irga, P. (2022). Urban green roofs promote metropolitan biodiversity: A comparative case study. Building and Environment, 207, 108458.

