## Supplementary materials for "Spatial factors predict variation in reports of human-wildlife interactions but not public attitudes towards a widespread urban carnivore, the red fox"

**Table S1.** Pairwise comparisons of the likelihood of reported bin-raiding foxes between UK regions.

| **Comparison** | **Z** | **p-value** |
| --- | --- | --- |
| London - East Midlands | 4.224 | **0.001** |
| London - East of England | 2.743 | 0.181 |
| London - North East | 3.568 | **0.016** |
| London - North West | 3.611 | **0.014** |
| London - Scotland | 4.992 | **<0.001** |
| London - South East | 3.703 | **0.010** |
| London - South West | 3.329 | **0.035** |
| London - Wales | 3.399 | **0.028** |
| London - West Midlands | 2.503 | 0.302 |
| London - Yorkshire and the Humber | 4.942 | **<0.001** |
| East Midlands - East of England | -1.792 | 0.786 |
| East Midlands - North East | 0.426 | 1 |
| East Midlands - North West | -1.204 | 0.982 |
| East Midlands - Scotland | 0.060 | 1 |
| East Midlands - South East | -1.477 | 0.928 |
| East Midlands - South West | -1.076 | 0.993 |
| East Midlands - Wales | -0.113 | 1 |
| East Midlands - West Midlands | -1.992 | 0.655 |
| East Midlands - Yorkshire and the Humber | 0.584 | 1 |
| East of England - North East | 1.808 | 0.776 |
| East of England - North West | 0.717 | 1 |
| East of England - Scotland | 2.113 | 0.568 |
| East of England - South East | 0.543 | 1 |
| East of England - South West | 0.732 | 1 |
| East of England - Wales | 1.406 | 0.947 |
| East of England - West Midlands | -0.230 | 1 |
| East of England - Yorkshire and the Humber | 2.459 | 0.329 |
| North East - North West | -1.364 | 0.957 |
| North East - Scotland | -0.407 | 1 |
| North East - South East | -1.553 | 0.902 |
| North East - South West | -1.270 | 0.974 |
| North East - Wales | -0.485 | 1 |
| North East - West Midlands | -1.959 | 0.678 |
| North East - Yorkshire and the Humber | 0.033 | 1 |
| North West - Scotland | 1.447 | 0.937 |
| North West - South East | -0.243 | 1 |
| North West - South West | 0.070 | 1 |
| North West - Wales | 0.898 | 0.998 |
| North West - West Midlands | -0.953 | 0.997 |
| North West - Yorkshire and the Humber | 1.879 | 0.732 |
| Scotland - South East | -1.797 | 0.783 |
| Scotland - South West | -1.276 | 0.973 |
| Scotland - Wales | -0.174 | 1 |
| Scotland - West Midlands | -2.335 | 0.410 |
| Scotland - Yorkshire and the Humber | 0.586 | 1 |
| South East - South West | 0.297 | 1 |
| South East - Wales | 1.107 | 0.991 |
| South East - West Midlands | -0.801 | 1 |
| South East - Yorkshire and the Humber | 2.192 | 0.510 |
| South West - Wales | 0.811 | 0.999 |
| South West - West Midlands | -0.950 | 0.997 |
| South West - Yorkshire and the Humber | 1.708 | 0.832 |
| Wales - West Midlands | -1.578 | 0.892 |
| Wales - Yorkshire and the Humber | 0.625 | 1 |
| West Midlands - Yorkshire and the Humber | 2.658 | 0.219 |

*Note.* A Type II ANOVA revealed that geographic region had a significant effect on reports of bin-raiding (χ²₁₀ = 49.707, p < 0.001). Tukey-adjusted post hoc comparisons between geographic regions are presented below with significant comparisons displayed in bold.

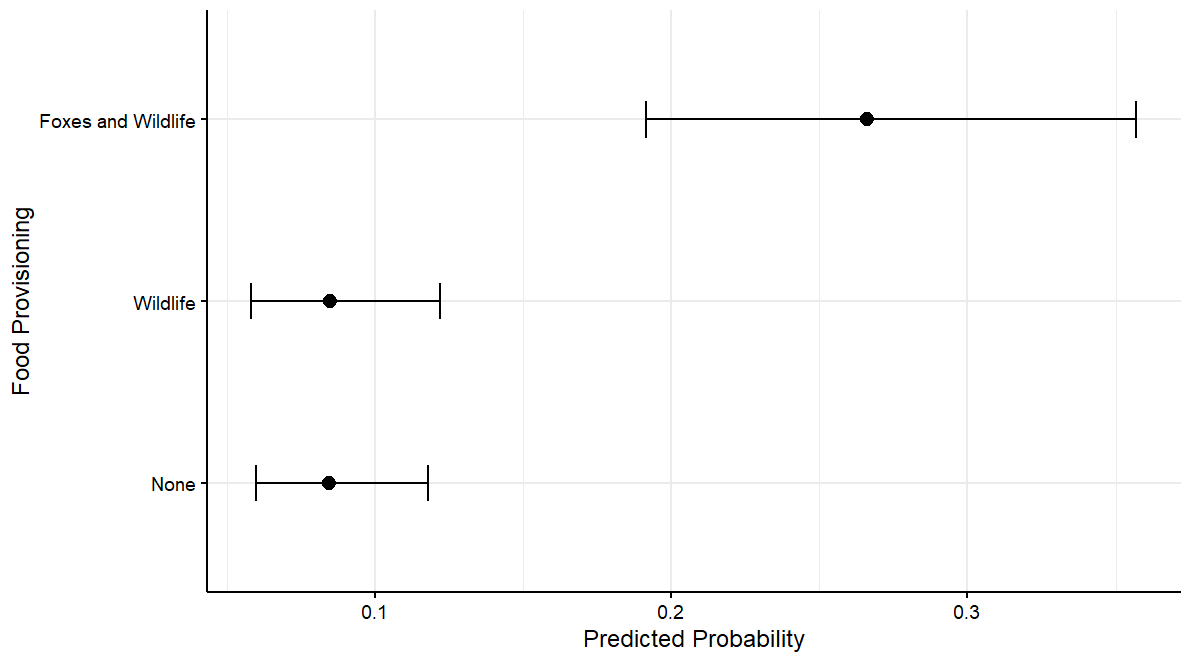

**Figure S1:** Predicted probabilities of people reporting the use of professional pest control services based on public wildlife feeding. Predictions are based on a logistic regression model controlling for geographic region, level of urbanisation, bin raiding experience, and attitudes towards foxes. Error bars represent 95% confidence intervals.

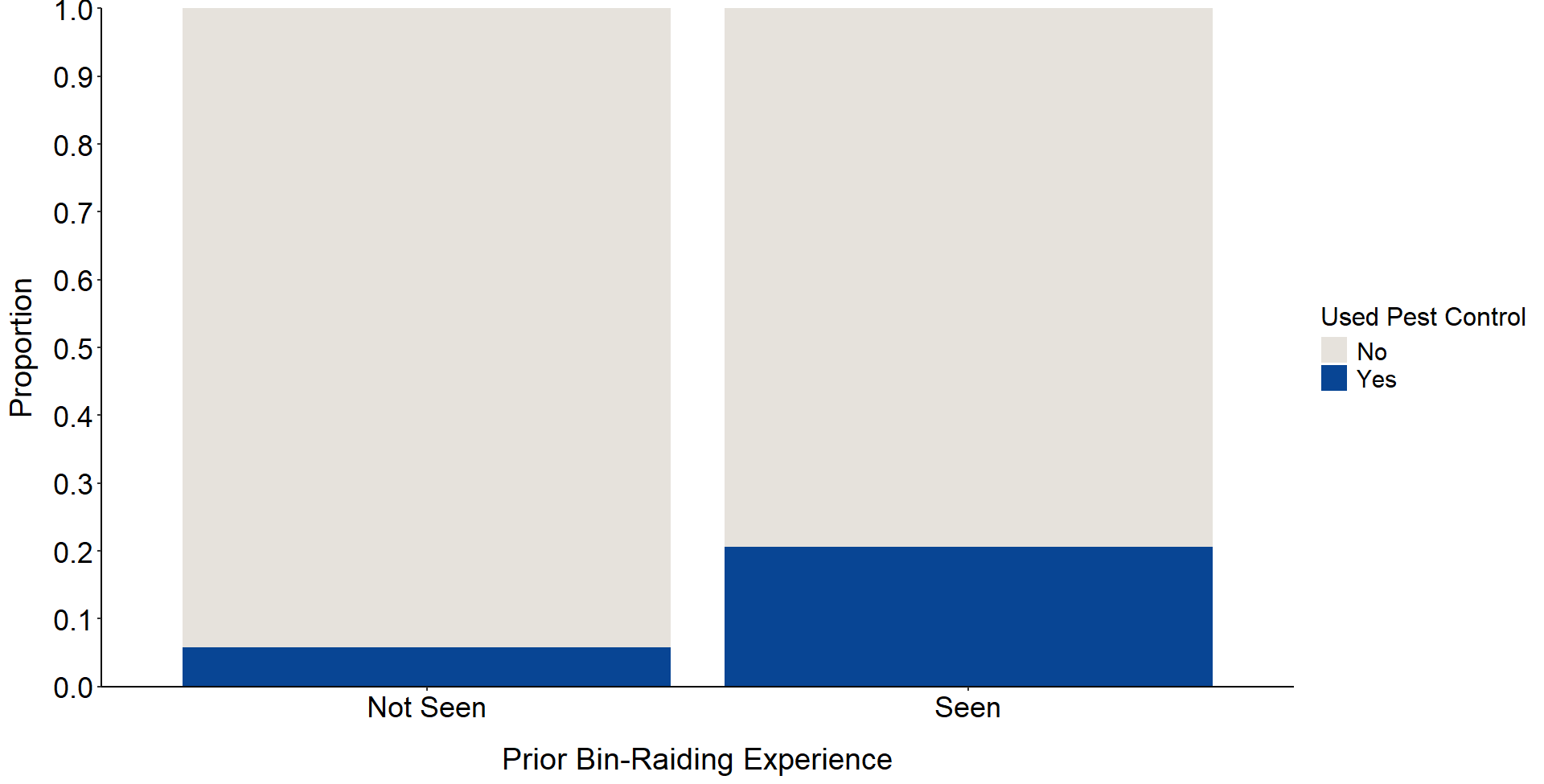

**Figure S2.** The proportion of people who reported having contacted professional pest control services to resolve a fox-related issue based on their reports of having (N = 277 participants) or not having (N = 998 participants) prior fox bin-raiding experience in their area.

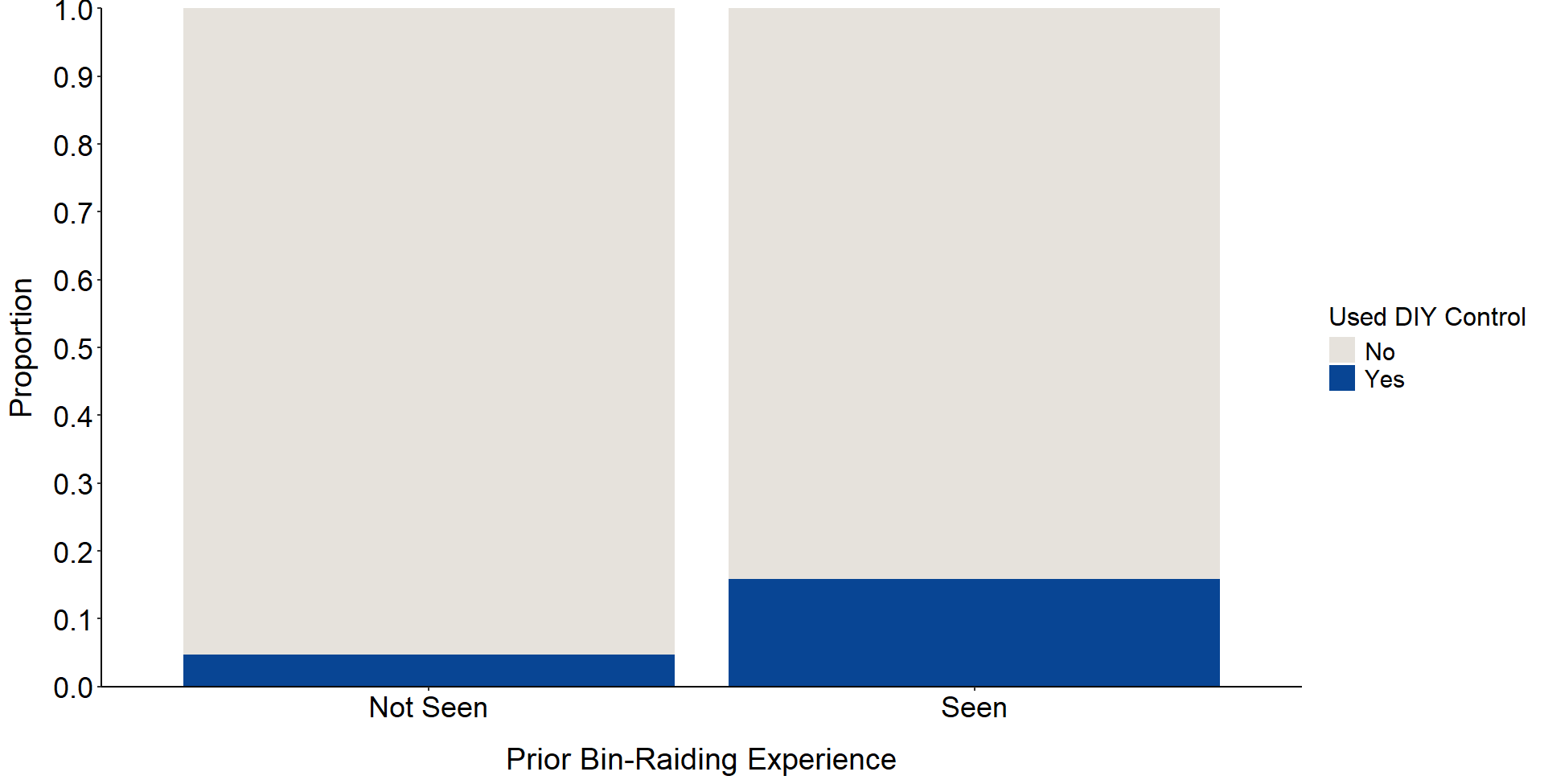

**Figure S3.** The proportion of people who reported using DIY control measures for foxes within their area based on their reports of having (N = 277 participants) or not having (N = 998 participants) seen a local fox raiding a bin.

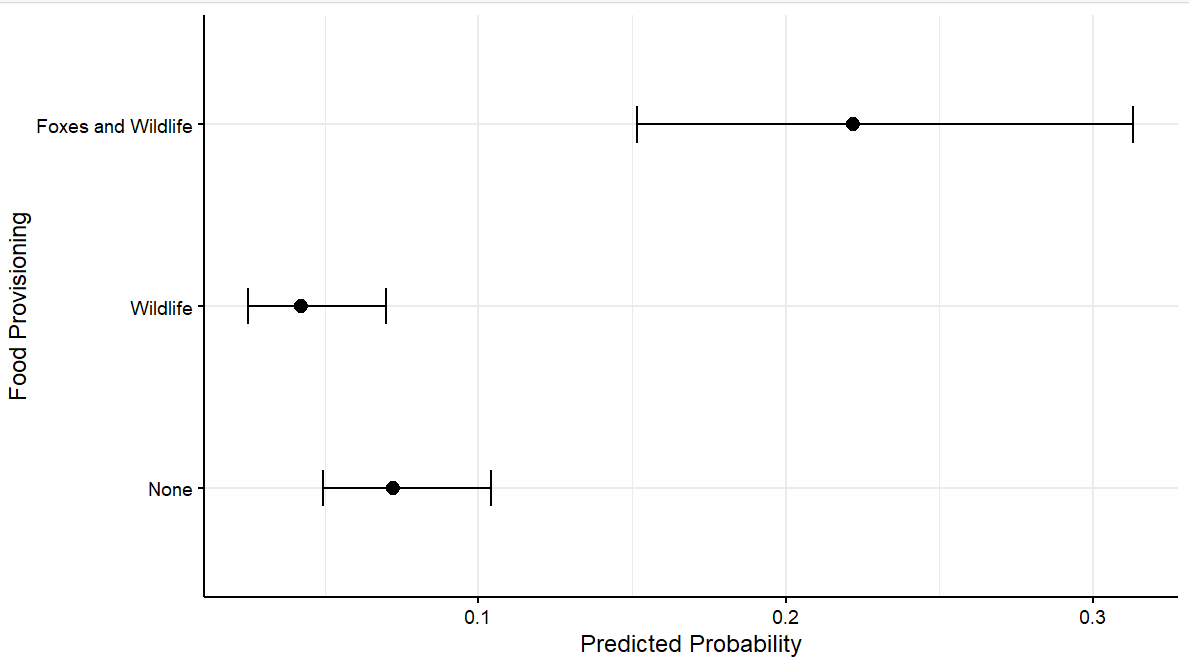

**Figure S4:** Predicted probabilities of people reporting the use of DIY fox control measures based on public wildlife feeding. Predictions are based on a logistic regression model controlling for geographic region, level of urbanisation, bin raiding experience, and attitudes towards foxes. Error bars represent 95% confidence intervals.

**Table S2.** Pairwise comparisons of the likelihood of the reported use of DIY fox control measures between UK geographic regions.

| **Comparison** | **Z** | **p-value** |
| --- | --- | --- |
| London - East Midlands | -2.965 | 0.104 |
| London - East of England | -1.283 | 0.972 |
| London - North East | -2.156 | 0.536 |
| London - North West | -0.565 | 1 |
| London – Scotland | -0.536 | 1 |
| London - South East | 0.270 | 1 |
| London - South West | -0.669 | 1 |
| London – Wales | -0.747 | 1 |
| London - West Midlands | 0.288 | 1 |
| London - Yorkshire and the Humber | -2.170 | 0.526 |
| East Midlands - East of England | 1.538 | 0.907 |
| East Midlands - North East | 0.204 | 1 |
| East Midlands - North West | 2.172 | 0.525 |
| East Midlands - Scotland | 2.155 | 0.537 |
| East Midlands - South East | 3.071 | 0.077 |
| East Midlands - South West | 1.846 | 0.752 |
| East Midlands - Wales | 1.479 | 0.927 |
| East Midlands - West Midlands | 2.661 | 0.218 |
| East Midlands - Yorkshire & Humber | 0.681 | 1 |
| East of England - North East | -1.043 | 0.994 |
| East of England - North West | 0.645 | 1 |
| East of England - Scotland | 1.462 | 1 |
| East of England - South East | 0.450 | 0.932 |
| East of England - South West | 0.260 | 1 |
| East of England - Wales | 0.260 | 1 |
| East of England - West Midlands | 1.318 | 0.966 |
| East of England - Yorkshire & Humber | -0.851 | 1 |
| North East - North West | 1.584 | 0.8894 |
| North East - Scotland | 1.569 | 0.896 |
| North East - South East | 2.281 | 0.447 |
| North East - South West | 1.355 | 0.959 |
| North East - Wales | 1.111 | 0.990 |
| North East - West Midlands | 2.093 | 0.583 |
| North East - Yorkshire & Humber | 0.341 | 1 |
| North West - Scotland | 0.006 | 1 |
| North West - South East | 0.777 | 1 |
| North West - South West | -0.150 | 1 |
| North West - Wales | -0.272 | 1 |
| North West - West Midlands | 0.731 | 1 |
| North West - Yorkshire & Humber | -1.487 | 0.924 |
| Scotland - South East | 0.753 | 1 |
| Scotland - South West | -0.153 | 1 |
| Scotland - Wales | -0.272 | 1 |
| Scotland - West Midlands | 0.708 | 1 |
| Scotland - Yorkshire & Humber | -1.480 | 0.927 |
| South East - South West | -0.880 | 0.999 |
| South East - Wales | -0.924 | 0.998 |
| South East - West Midlands | 0.052 | 1 |
| South East - Yorkshire & Humber | -2.334 | 0.4106 |
| South West - Wales | -0.132 | 1 |
| South West - West Midlands | 0.826 | 0.999 |
| South West - Yorkshire & Humber | -1.223 | 1 |
| Wales - West Midlands | 0.883 | 0.999 |
| Wales - Yorkshire & Humber | -0.935 | 0.998 |
| West Midlands - Yorkshire & Humber | 0.883 | 0.611 |

*Note*. A Type II ANOVA revealed that geographic region had a significant effect on the reported use of fox control measures (χ²₁₀ = 19.332, p = 0.036). Tukey-adjusted post hoc comparisons between geographic regions are presented, however, none of the comparisons were significant.
